# Maternal SSRI treatment during offspring development results in long-term behavioral, cellular, and neuroimaging disruptions

**DOI:** 10.1101/205708

**Authors:** Susan E. Maloney, Rachel Rahn, Shyam Akula, Michael A. Rieger, Katherine B. McCullough, Christine Jakes, Selma Avdagic, Krystal Chandler, Amy L. Bauernfeind, Joseph P. Culver, Joseph D. Dougherty

## Abstract

Serotonergic dysregulation is implicated in psychiatric disorders, including autism spectrum disorders (ASD). Epidemiological studies suggest selective serotonin reuptake inhibitor (SSRI) treatment during pregnancy may increase ASD risk in offspring, however it is unclear from these studies whether ASD susceptibility is related to the maternal diagnosis or if treatment poses additional risk. Here, we exposed mouse dams to fluoxetine and characterized the offspring to isolate possible effects of SSRI exposure on ASD-relevant behaviors. We demonstrate social communication and interaction deficits and repetitive behaviors, with corresponding dendritic morphology changes in pertinent brain regions. Also, using a novel application of optical intrinsic signal imaging, we show altered stimulus-evoked cortical response and region-specific decreases in functional connectivity. These findings indicate drug exposure alone is sufficient to induce long-term behavioral, cellular, and hemodynamic-response disruptions in offspring, thus contributing to our understanding of ASD pathogenesis, risk and mechanism, as well as the developmental role of serotonin.

## Introduction

Dysregulation of the serotonin (5HT) system is implicated in numerous psychiatric disorders (Nordquist & Oreland, 2010). This system innervates the entire CNS, allowing 5HT to influence a variety of behavioral functions (Smythies, 2005). During prenatal development, 5HT is one of the earliest neuromodulators to become active, and levels of 5HT, the expression of the 5HT transporter and receptors are at their peak, allowing 5HT to play widespread trophic roles in brain development (Whitaker-Azmitia, 2010). The placenta is a transient source of 5HT to the fetal forebrain, and perturbations in placental 5HT output can alter neurodevelopment (Goeden et al., 2016). This suggests that altering fetal-or maternally-derived 5HT can impact circuit development, possibly increasing risk for psychiatric disorders.

5HT dysregulation is implicated in neurodevelopmental disorders such as autism spectrum disorder (ASD): ~25% of ASD patients exhibit elevated 5HT levels in whole blood platelets (Benza & Chugani, 2015); changes to 5HT can either worsen or alleviate certain symptoms (Hollander et al., 2005; McDougle et al., 1993, 1996); increased 5HT axons are observed postmortem (Azmitia, Singh, & Whitaker-Azmitia, 2011), and PET studies demonstrate altered 5HT synthesis *in vivo* (Chugani et al., 1997, 1999). Thus, genes or environmental agents that dysregulate 5HT are potential risk factors for ASD.

5HT is a dominant target for treatment in many psychiatric conditions through frequently prescribed medications such as selective serotonin reuptake inhibitors (SSRIs). SSRIs are highly prescribed in pregnant women for mood disorders with prevalence of 5-13% (Cooper, Willy, Pont, & Ray, 2007). A recently published meta-analysis reported a significant case-control association between maternal antidepressant use and ASD risk in offspring, which remained when adjusted for maternal psychiatric history (Mezzacappa et al., 2017). Meta-analyzed cohort studies replicated the SSRI-ASD link, but not the independence from psychiatric history. Thus, the human studies remain controversial. In addition, direct causality and biological mechanisms cannot be inferred from epidemiological studies. Animal studies can provide clear indication as to whether transient SSRI exposure can alter long-term behaviors in mammals, independent of maternal psychiatric stress, and provide ready access to the underlying neurobiology. We therefore developed a rodent model of maternal SSRI exposure, in the absence of maternal stress, to determine if drug alone induces behavioral disruptions related to the core symptoms of ASD in offspring, and conducted a parallel analysis of cellular morphology in cortical regions mediating these behaviors.

Many studies have attempted to identify functional neuroimaging correlates of ASD in humans but also with mixed results (Hull et al., 2017). These studies are challenged by the variety of factors contributing to ASD risk, as patients may have different underlying etiologies and thus corresponding heterogeneity in neuroimaging phenotypes. A mouse system for examination of mesoscale functional connectivity would allow for well-powered analyses of a single risk factor, such as SSRI exposure. Therefore, we employed an optical intrinsic signal (OIS) imaging pipeline and provided the first description of alterations in functional connectivity and evoked responses in a mouse model of ASD environmental risk.

Overall, across multiple exposure durations, we found strong evidence supporting the hypothesis that transient exposure to SSRIs has long-term consequences on behavior, anatomy, and functional connectivity in the mammalian brain.

## Results

### Development of SSRI maternal exposure models

To determine the potential of maternal SSRI exposure to induce behavioral disruptions in offspring relative to ASD symptomatology, we exposed mouse dams to fluoxetine (FLX) during gestation and lactation and examined offspring behaviors during development, the juvenile stage, and adulthood (**Figure 1A, Table S1**). In most countries, FLX was the first SSRI to become available for clinical use (Hiemke & Hartter, 2000); therefore, FLX is likely to be the most-represented antidepressant in the epidemiology studies of SSRI use during pregnancy and ASD risk. As genetic factors are clearly an important causation of ASD (Geschwind, 2008), it is likely that environmental contributions to ASD risk interact with existing genetic susceptibility (Hertz-Picciotto et al., 2006; Klei et al., 2012). It has been suggested that environmental factors that might modulate social behavior or language could tip the balance towards ASD in children with genetic vulnerability (Geschwind, 2008). As we initially thought SSRI exposure alone might be a relatively modest factor, we exposed *Celf6* mutant mice, which exhibit a partial ASD-like phenotype (Dougherty et al., 2013), to maternal FLX and analyzed offspring behavior for possible potentiation of the ASD-like phenotype (*Cef6*-Extended). Further, *Celf6* is enriched in 5HT-producing cells and when deleted results in a decrease in brain 5HT levels. Thus, we hypothesized that early exposure to FLX in *Celf6* mutant mice may interact synergistically on the 5HT system.

**Figure 1.**
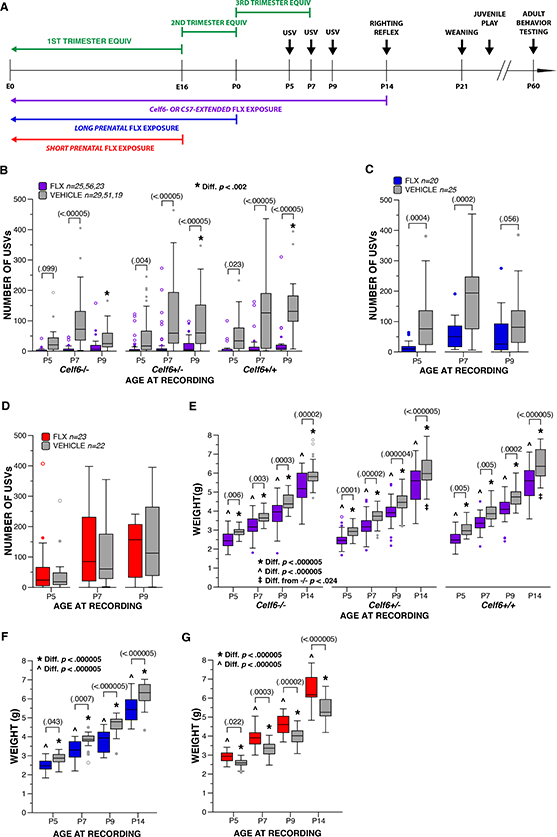
Maternal FLX throughout pregnancy alters early communicative behavior. (A) Schematic of paradigm for maternal FLX, with approximate equivalents in brain development to human pregnancy, and mouse age for each behavioral test. (B-D) Boxplots of number of USVs at P5, P7 and P9 from *Celf6*-Extended (Drug, *p*<.000005; Age × Drug × Genotype Interaction, *p*=.049), Long Prenatal (Drug, *p*=.0001) and Short Prenatal (Drug, *p*>.05) FLX and VEH mice (thick horizontal lines signify respective group medians, boxes are 25^th^ – 75^th^ percentiles, whiskers are 1.5 × IQR, closed and open circles depict outliers). (E-G) Boxplot of weight (g) at P5, P7, P9 and P14 of *Celf6*-Extended (Drug, *p*<.000005), Long Prenatal (Drug, *p*=.00004) and Short Prenatal (Drug, *p*=.000008) FLX and VEH *Cefl6* mutant and WT littermates All mice gained weight with age.

We also examined different pre- and postnatal durations of FLX to establish periods of vulnerability. Epidemiological studies are inconsistent regarding the trimesters of pregnancy most vulnerable to SSRI-induced ASD risk. To address this, we used three FLX durations, corresponding to periods of brain development approximating the trimesters of human pregnancy. Our designation of “Extended FLX” corresponded to the entire duration of the pregnancy and a recommended period of nursing (1 year) in humans (embryonic day (E)0, through postnatal day [P]14) (Dobbing & Sands, 1979; Levitt, 2003). Both *Celf6* and C57BL/6J mice were exposed for this duration (Celf6-Extended and C57-Extended). “Long Prenatal” (E0-P0) exposure approximated the first and second trimesters of human pregnancy. “Short Prenatal” (E0-E16) approximated the first trimester of human pregnancy (**Figure 1A**). Only C57BL/6J mice were used for Prenatal-only exposures. Overall, our experimental design enabled analysis of both gene × environment interaction and exposure duration effects on behaviors relevant to the symptoms of ASD.

### Maternal FLX disrupts early communicative behavior in pup offspring

We examined developmental behavior, physical milestones and reflexes in our FLX mice. Isolation-induced pup ultrasonic vocalizations (USVs) are considered a strongly conserved affective and communicative display that elicits maternal behaviors (Hofer, Shair, & Brunelli, 2002). This behavior has a distinct developmental trajectory, allowing its use for the study of both early communication and neurobehavioral development in infant rodents (Igor Branchi, Santucci, & Alleva, 2001). At P5, 7, and 9, we observed robust decreases in USVs when FLX lasted through pregnancy. No influence of sex was observed for developmental analyses, therefore all data reported below are collapsed for sex. Output from statistical tests is fully reported in **Table 1**. Specifically, *Celf6*-Extended exposure to FLX reduced USVs independent of *Celf6* genotype (*p*<.000005, **Figure 1B**). *Celf6* mutation also reduced USVs in vehicle (VEH)-exposed pups (*p*=.049), replicating previous work (Dougherty et al., 2013). Post hoc tests revealed FLX-induced USV reduction at each age across all mice (*p*<.024), except for P5 and P7 *Celf6*^−/−^ mice when USVs were already low. USV calls from FLX pups also were altered in spectral and temporal features, including average pitch, percent calls containing a pitch jump, and pitch range of calls containing a pitch jump (*p*<.020; **Figure S1A-C**). Robust reductions in the duration time of calls and the pitch range of flat calls were observed in FLX pups (*p*<.000005; **Figure S1D,E**). *Celf6* mutation did not influence spectral or temporal features of USVs alone or through an interaction with extended FLX.

**Table 1:**
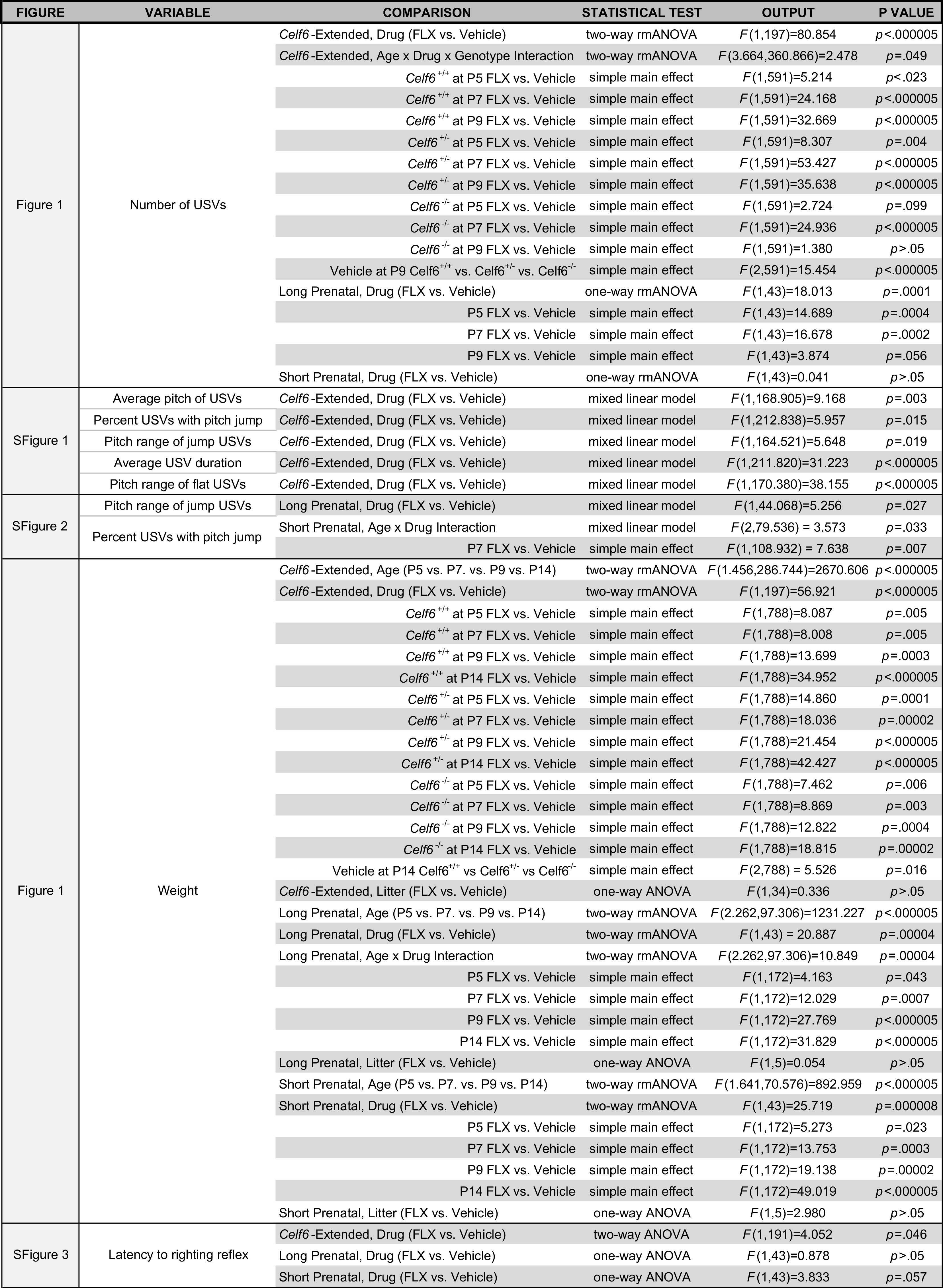
Statistical summary for Figures 1, S1-3.

Since the impact of FLX alone was so strong, and independent of *Celf6* mutation in the Celf6-Extended cohort, we examined the impact of prenatal-only exposure to FLX on USV in C57BL/6J mice. Long Prenatal exposure to FLX also reduced USVs (*p*=.0001; **Figure 1C**). This FLX-induced reduction occurred at P5 and P7 (*p*<.0005), with a trend at P9 (*p*=.056). Examination of spectral and temporal features showed Long Prenatal exposure only altered the pitch range of flat calls (*p*=.027; **Figure S2A-D**). Short Prenatal exposure to FLX did not influence pup USV production (**Figure 1D**). However, at P7, FLX pups produced a greater percent of calls containing a frequency jump (*p*=.007; **Figure S2F-J**), without altering other features. Taken together, these findings suggest FLX, when continued through pregnancy, induced early communicative deficits in mice in the form of USV reductions, yet FLX limited to early pregnancy did not influence production rate. Further, the effect of FLX was so robust that additional interaction with *Celf6* mutation on USVs was not observed.

### Developmental assessment of physical milestones and reflexes

USV suppression may be a consequence of perturbation of specific CNS circuits due to FLX exposure. However, an alternative explanation is that USV is suppressed by a FLX-induced gross developmental delay. To explore this possibility, we examined other developmental traits of FLX pups. As a measure of general health, we compared the weight of FLX and VEH mice on P5, 7, 9 and 14. Mice in all cohorts increased in weight across developmental time points, as expected (*p*<.000005, **Figure 1E-G**), yet the duration of FLX exposure influenced weight. All *Celf6*-Extended and Long Prenatal FLX mice weighed less than VEH pups (*p*<.00005, **Figure 1E,F**) regardless of genotype at each age examined (*p*<.044). Interestingly, Short Prenatal FLX resulted in increased weight compared to VEH (*p*=.000008, **Figure 1G**) at all ages examined (*p*<.023). Weight differences were not due to differences in litter sizes produced by FLX and VEH dams, as litter size differences were not detected for any cohort (*p*>.05). Further assessment of developmental milestones revealed that FLX exposure had no effect on the timing of pinna detachment (by P5) or eye opening (by P14; data not shown). To assess early gross locomotor abilities and to evaluate general body strength, we examined righting reflex at P14. When collapsed across genotypes, FLX pups in the *Celf6*-Extended cohort exhibited a longer latency to right compared to VEH pups (*p*=.046; **Figure S3A**). No difference in latency to right was observed in the Long Prenatal cohort (**Figure S3B**), and in the Short Prenatal cohort, a strong trend towards increased time to right was observed in FLX pups compared to VEH pups (*p*=.057; **Figure S3C**). The developmental data indicate age-appropriate physical milestones were achieved, indicating FLX did not induce gross developmental delay; however, developmental reflexes were minimally influenced by FLX and weight was affected across development suggesting FLX exposure did induce marginal developmental perturbation in pups.

### Maternal FLX disrupts adult social behaviors

Deficits in social communication and social interaction are varied among autistic individuals, and include failure to initiate or respond to social interaction, abnormal social approach, and difficulties adjusting behavior to suit various social contexts (American Psychiatric Association, 2013). Therefore, we tested our mice in multiple social behavior assays, each designed to assess a distinct aspect of social behavior. The full-contact juvenile play assay was used to assess social interaction behaviors in FLX mice, and in adulthood, we examined social approach behaviors and possible disruptions to behaviors in the specific context of social dominance hierarchies.

Maternal FLX exposure disrupted social approach and specific social hierarchy behaviors in adulthood, but not juvenile social interactions. The social approach task was designed to examine sociability, or the typical preference of spending more time investigating a novel conspecific (social stimulus) versus a novel object (empty cup stimulus) (Moy et al., 2004). First, during the habituation trial, no chamber side bias was observed for any cohort (**Figure S4A-D**). However, during test trials we observed decreased sociability in the Celf6-Extended and Long Prenatal exposure groups. Output from statistical tests is fully reported in **Table 2**. Specifically, in the Celf6-Extended exposure group, female FLX *Celf6*^+/+^ mice (*p*=.038) failed to display a preference for the social stimulus (*p*>.05; VEH females, *p*=.005; **Figure 2A**) and spent less time investigating the social stimulus compared to VEH *Celf6*^+/+^ females (*p*=.050). When collapsed across sex, VEH mice spent more time compared to FLX mice investigating both stimuli overall (*p*=.020). Yet, the typical preference for social stimulus was observed for all FLX and VEH mice, with comparable time spent investigating the social stimulus (*p*<.022). As *Celf6* mutation did not potentiate the impact of FLX, we continued our examination of social approach behaviors without manipulation of *Celf6* genotype for the Long and Short Prenatal cohorts. Long prenatal exposure resulted in disruptions to sociability regardless of sex (*p*=.0004). FLX mice failed to display a preference for the social stimulus (*p*>.05; VEH, *p*<.000005; **Figure 2B**), and spent significantly less time investigating the social stimulus compared to VEH mice (*p*=.0001, **Figure 2B**). Short prenatal exposure did not disrupt sociability: both FLX and VEH spent more time investigating the social stimulus than the empty cup (FLX, *p*=.001; VEH, *p*=.001; **Figure 2C**), and a similar time was spent investigating the social stimulus by both groups. Finally, during the preference for social novelty trial, more time was spent investigating the novel mouse compared to the familiar mouse in all cohorts (*p*<.045; **Figure S4A-D**). Comparable activity levels were detected for all groups in this task (**Figure S4E-G**), ruling out hypoactivity as a confound. Taken together, these data indicate maternal FLX influenced sociability only when continued throughout pregnancy. FLX exposure limited to early pregnancy did not significantly influence sociability in our mice.

**Table 2:**
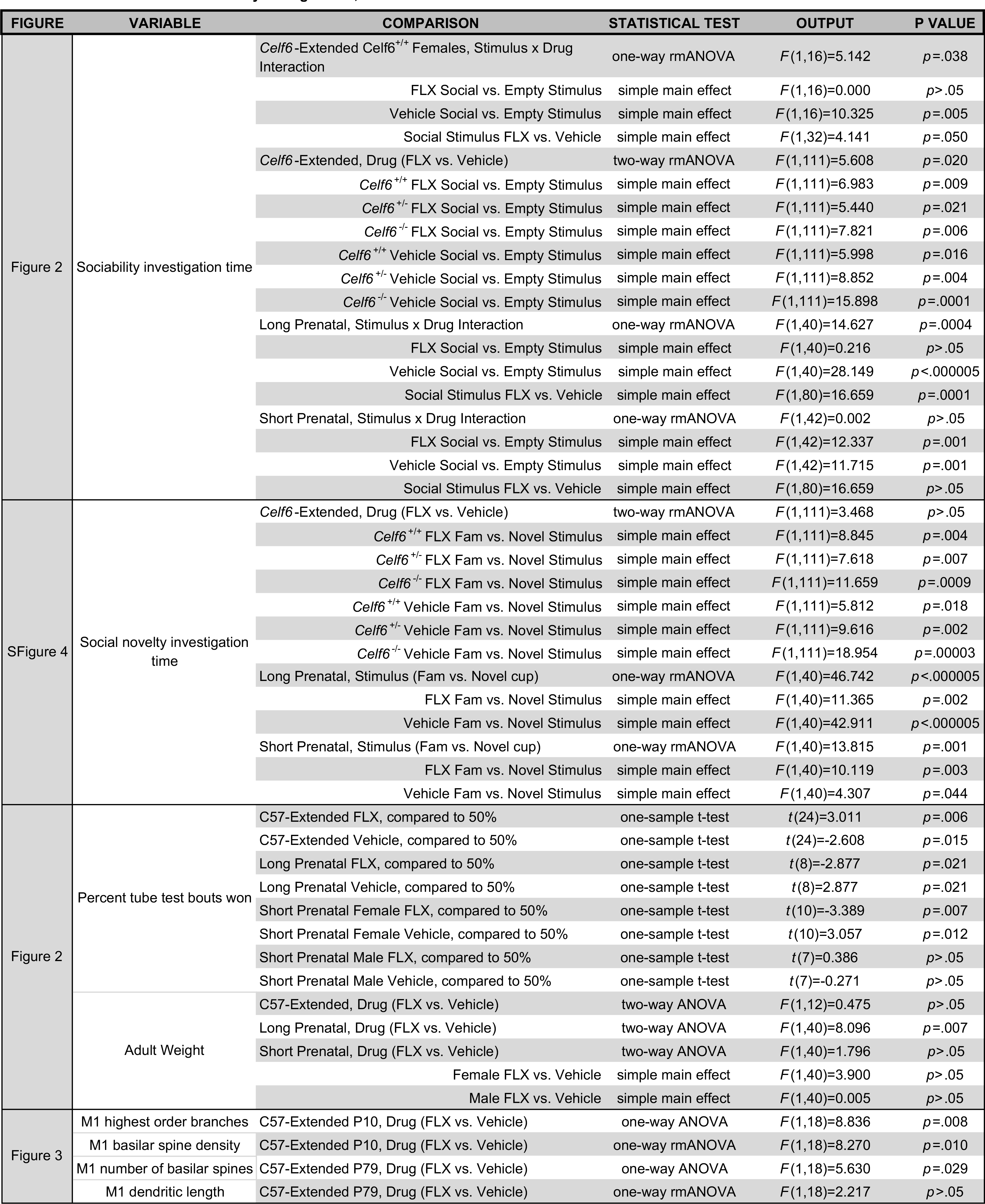
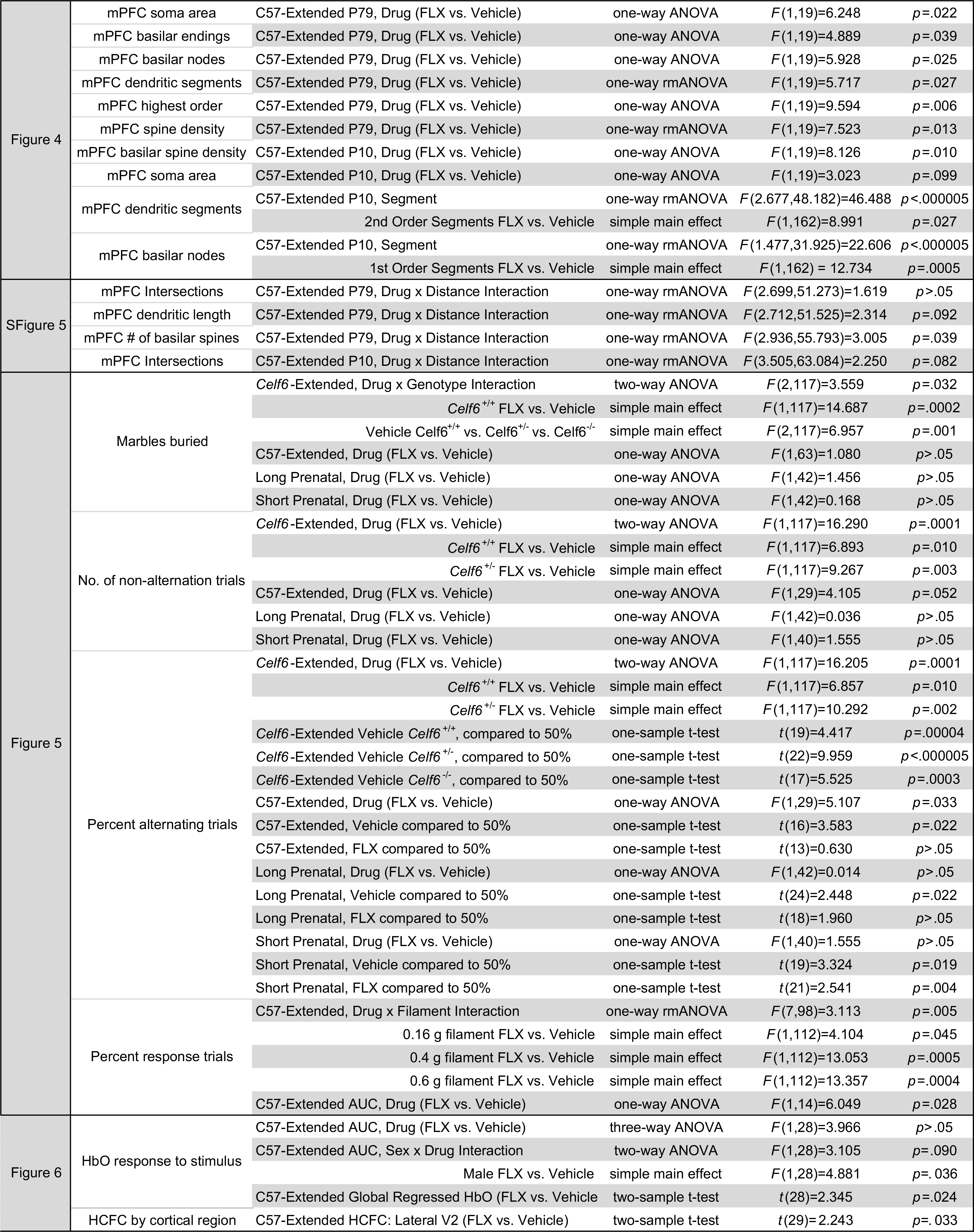
Statistical summary for Figure 2-5, S4-5.

**Figure 2.**
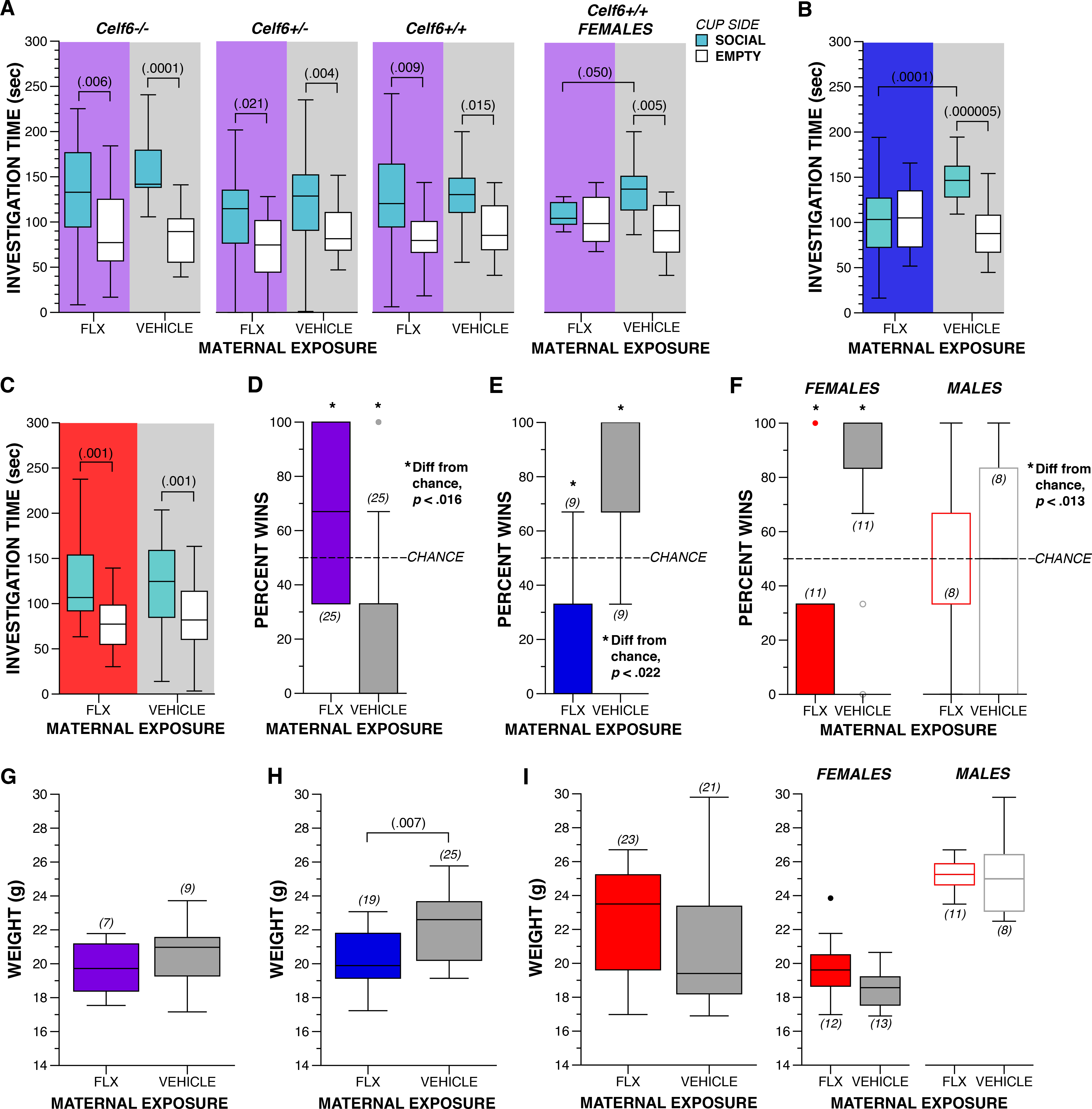
Maternal FLX disrupts adult sociability and social dominance hierarchy behaviors. (A-C) Boxplot of time spent investigating social and empty cups during social approach test by *Celf6*-Extended (Drug, *p*=.020; Female *Celf6*^+/+^ mice Stimulus × Drug, *p*=.038), Long Prenatal (Stimulus × Drug, *p*=.0004) and Short Prenatal (Stimulus × Drug, *p* > .05) FLX and VEH mice. (D-F) Boxplot of percent wins during tube test of social dominance during between C57-Extended, Long Prenatal, and Short Prenatal FLX and VEH adult mice. (G-I) Boxplot of weight of C57-Extended, Long Prenatal, and Short Prenatal FLX and VEH adult mice.

As only *Celf6* wild type (WT) mice in the *Celf6*-Extended cohort displayed sociability deficits, we chose to examine full-contact social behaviors in C57BL/6J juveniles in a separate C57-Extended cohort. We did not observe abnormal social interactions in these mice in the juvenile play assay. Specifically, FLX and VEH mice exhibited a comparable number and duration of anogenital and head-to-head sniffing, and sniffing behaviors directed toward FLX and VEH mice by the stimulus partners were also similar (data not shown). Unlike the social approach task, we did not observe altered social behaviors in the juvenile play assay. However, in social approach only the FLX mouse has control over timing and duration of interactions, while in juvenile play deficits in social behaviors with FLX treatment could be masked because interactions were also initiated by the unexposed stimulus mouse.

Finally, we examined social hierarchy behaviors in our mice to determine if maternal FLX exposure influences behavior in this specific social context. Groups of mice display social hierarchies with dominant and submissive group members (Hayashi, 1993), and we assessed this using the tube test for social dominance. For this task, sex-matched mice from different experimental groups are directly compared. Due to the complexity of experimental groups in the *Cef6*-Extended cohort, we only examined tube test behavior between FLX and VEH mice in the C57-Extended cohort. We observed an interesting influence of FLX duration on dominance. C57-Extended FLX resulted in increased dominant behavior (**Figure 2D**, FLX wins greater than by chance, *p*=.006; VEH wins fewer than by chance, *p*=.015). In contrast, both maternal FLX cohorts restricted to prenatal development induced submissive behaviors in adulthood: Long Prenatal FLX resulted in fewer wins relative to chance (*p*=.021; **Figure 2E**). Short Prenatal exposure influenced dominance behavior only in female mice, which again won fewer bouts than expected by chance (*p*=.007; **Figure 2F**). No difference was observed for FLX or VEH males in the Short Prenatal cohort. These alterations in dominance were not due to differences in animal size between drug exposure groups as adult weights did not correspond to increased dominance in a simple way. Specifically, at available power we did not detect differences in adult weight in C57-Extended FLX mice (*p*>.05; **Figure 2G**). Long Prenatal FLX resulted in a decrease in weight compared to VEH, that was independent of sex (*p*=.007; **Figure 2H**) and dominance performance. We did not detect a weight difference in the males or females of the Short Prenatal cohort (**Figure 2I**), despite altered sex-specific dominance. Taken together, these data suggest perinatal FLX exposure via the mother influences social behaviors during adulthood, long after drug exposure occurred, with specific disruptions to sociability and behavior in the specific context of dominance. Further, prenatal versus postnatal exposure may differentially influence behavioral circuits underlying dominance behaviors.

### Maternal FLX results in persistent changes to layer V pyramidal cell dendrite morphology

We next examined the neuronal morphologies of mice with the most consequential exposure condition (C57-Extended) to identify long-term cellular changes induced by FLX exposure within key cortical regions (**Figure 3A**). ASD is known to be a disorder of the synapse and information processing (Won, Mah, & Kim, 2013) and altered dendritic spine densities have been observed in postmortem ASD cortical tissues, suggesting changes to excitatory synapse numbers (Hutsler & Zhang, 2010). Thus we hypothesized that dendritic morphology and synaptic densities of neurons in areas of the mouse brain relevant to the social behavior deficits would also be altered.

**Figure 3.**
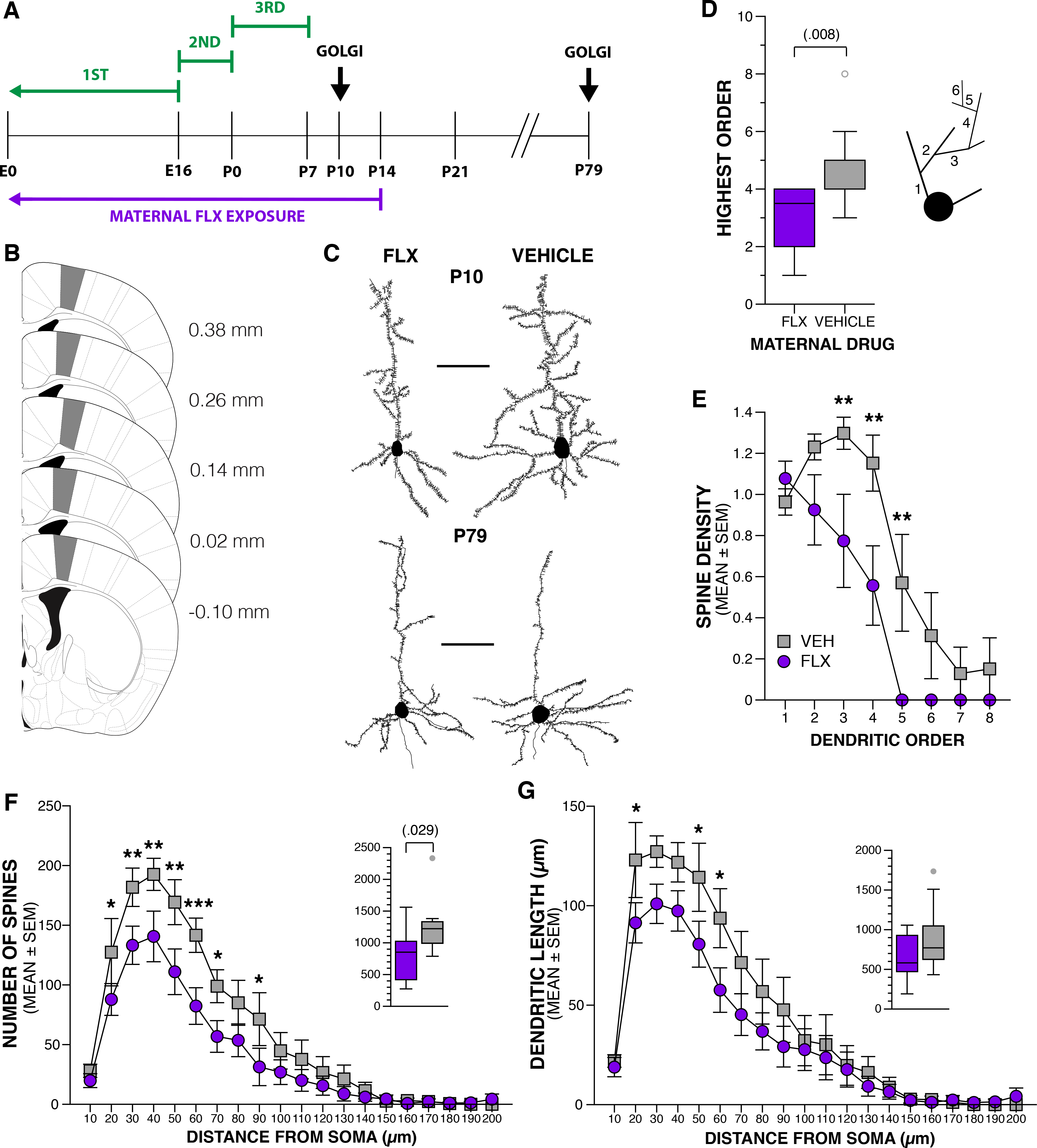
Maternal FLX alters dendritic morphology of M1 layer V pyramidal neurons. (A) Schematic of the experimental paradigm. (B) M1 area used for neuron selection marked in grey. Numbers indicate distance from bregma. (C) Representative tracings of Golgi-Cox impregnated M1 layer V neurons from P10 and P79 FLX and VEH brains. Scale bars are 50 μm. (D) Boxplot of highest branch order of basilar dendrites of layer V neurons from FLX and VEH brains at P10 (n=10 neurons from 5 FLX mice and n=10 neurons from 5 VEH mice). Inset: schematic of dendritic branch orders, where 1^st^ order branches initiate at the soma, and each subsequent branch order extends from the previous, lower order. (E) Spine density per basilar dendritic branch order in neurons from FLX and VEH P10 brains (data are means ± SEM; Drug, *p*=.010). (F) Number of basilar dendritic spines per 10 μm distance from the soma in neurons from FLX and VEH P79 brains (n=10 neurons from 5 FLX mice and n=10 neurons from 4 VEH mice;). Inset boxplot of total number of dendritic spines per drug group. (G) Basilar dendritic length per 10 |im distance from the soma in neurons from FLX and VEH P79 brains. Inset boxplot of total dendritic length per drug group. Statistical significance, \*\*\**p*<.001, \*\**p*<.01, \**p*<.05.

Following the robust decrease in USVs at P5-P9 in FLX mice, we sought to examine whether there were corresponding morphological changes in layer V pyramidal neurons in primary motor cortex (M1; **Figure 3B**). The specific portion of M1 sampled corresponded to the area which sends projections to the nucleus ambiguus, the motor nucleus of cranial nerves IX and X that innervates the muscles of the soft palate, pharynx, and larynx, that is active during adult mouse USV (Arriaga, Zhou, & Jarvis, 2012). We quantified the dendritic morphology of these neurons (**Figure 3C**) at P10. Output from statistical tests for this section is fully reported in **Table 2**. A clear decrease in extent of basilar dendrite branching was observed in neurons from FLX brains (*p*=.008; **Figure 3D**). In addition, among the 3^rd^-5^th^ branch order segments, FLX reduced the spine density (*p*=.010; **Figure 3E**). We also quantified dendritic features of these same neurons in adult P79 mouse brains to determine if alterations to morphology are transient or persistent (**Figure 3C**). We observed a decrease in basilar dendritic spines in FLX neurons (*p*=.029; **Figure 3F**), with the largest reduction between 30–60 μm from the soma (*p*<.004). Spine density remained similar between the FLX and VEH groups due to a concurrent, although statistically non-significant, reduction in branch length in FLX neurons (*p*>.05; **Figure 3G**). Extent of branching was no longer different between drug groups at P79. Total dendritic length, dendritic field, soma area and aspects of dendritic complexity were indistinguishable between the two drug groups at P10 and P79 (**Table S2**). These findings corroborate previous reports of reduced basilar dendritic spines on layer V neurons of the M1 in adult rats exposed to FLX from P0–P4 that exhibited poor motor performance on the rotarod task (Lee & Lee, 2012). In addition, these data reveal the novel finding that developmental exposure to maternal FLX coincides with a reduction in basilar dendritic branching and spine density and is correlated with a dramatic reduction in USV production.

C57-Extended FLX resulted in increased dominance in adulthood (**Figure 2D**) and previous research showed excitatory synaptic efficacy of layer V pyramidal cells in the medial prefrontal cortex (mPFC) regulates social dominance rank in the mouse (Wang et al., 2011). Therefore, to determine if perinatal FLX exposure altered features of these neurons, we quantified their dendritic morphology at P79 (**Figure 4A,B**). FLX resulted in smaller soma area (*p*=.022; **Figure 4C**) and less complex basilar dendritic branching. Specifically, the numbers of basilar endings (*p*=.039; **Figure 4D**) and nodes (*p*=.025; **Figure 4E**) were reduced. Dendritic branch orders displayed fewer overall dendritic segments *p*=.027; **Figure 4F**), particularly 2^nd^ order segments (*p*=.003). Neurons from FLX brains also had less extensive branching (*p*=.006; **Figure 4G**), and spine density was reduced in higher-order branches of FLX neurons (*p*=.013; **Figure 4H**), specifically 4^th^ and 5^th^ order branches (*p*<.01). Similar results were found by applying the Sholl method to spatially analyze aspects of dendritic complexity (**Figure S5A-C**).

**Figure 4.**
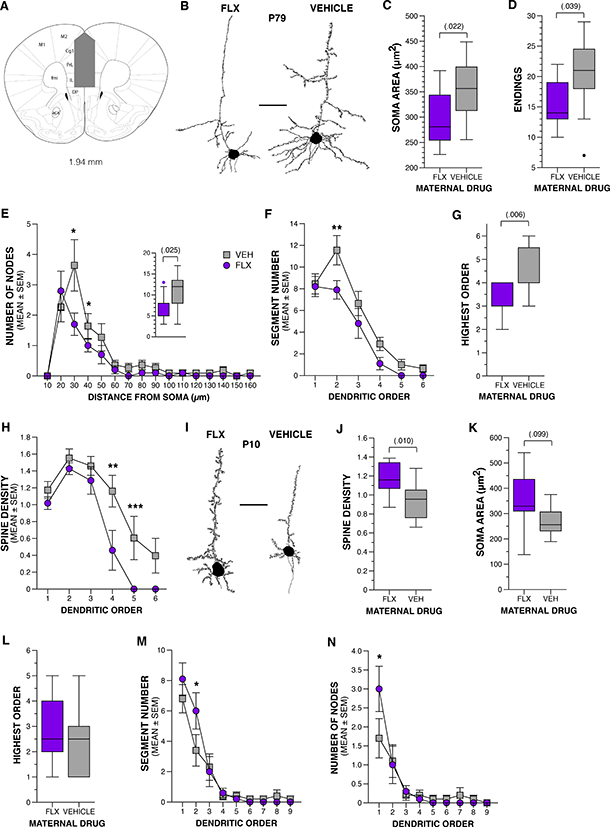
Maternal FLX alters dendritic morphology of mPFC layer V pyramidal neurons. (A) mPFC area used for neuron selection marked in grey. Numbers indicate distance from bregma. (B) Representative tracings of Golgi-Cox impregnated mPFC layer V neurons from P79 FLX and VEH brains. Scale bar is 50 μm. (C) Boxplot of soma area for neurons from FLX and VEH brains at P79 (n=10 neurons from 6 FLX mice and n=11 neurons from 6 VEH mice). (D) Boxplot of number of basilar dendritic endings of neurons from FLX and VEH brains at P79. (E) Number of branching nodes per 10 μm distance from the soma neurons from FLX and VEH P79 brains (data are means ± SEM). Inset boxplot of total number of branching nodes per drug group. (F) Number of branch segments per basilar dendritic branch order in layer V neurons from FLX and VEH P79 brains (Drug, *p*=.027). (G) Boxplot of highest order branching of basilar dendrites of layer V neurons from FLX and VEH brains at P79. (H) Spine density per basilar dendritic branch order in layer V neurons from FLX and VEH P79 brains. (I) Representative tracings of Golgi-Cox impregnated mPFC layer V neurons from P10 FLX and VEH brains. Scale bar is 50 μm. (J) Boxplot of spine density on basilar dendrites of layer V neurons from FLX and VEH brains at P79. (K) Boxplot of soma area for neurons from FLX and VEH brains at P10 (n=10 neurons from 5 FLX mice and n=10 neurons from 5 VEH mice). (L) Boxplot of highest order branching of basilar dendrites of layer V neurons from FLX and VEH brains at P10. (M) Number of branch segments per basilar dendritic branch order in neurons from FLX and VEH P10 brains (Drug, *p*>.05; Bonferroni corrected). (N) Number of branching nodes per basilar dendritic branch order in neurons from FLX and VEH P10 brains (Drug, *p*>.05, Bonferroni corrected). Statistical significance, \*\*\**p*<.001, \*\**p*<.01, \**p*<.05.

To determine if these adult deficits were apparent in early postnatal ages, we examined these neurons from P10 mice exposed to FLX or VEH (**Figure 4A,I**). Interestingly, the findings are in the opposite direction from that observed in P79 brains. We observed an increase in basilar dendritic spine density in neurons exposed to FLX (*p*=.010; **Figure 4J**), and FLX neurons had a larger soma size compared to VEH neurons, although this did not reach statistical significance (*p*=.099; **Figure 4K**). They also had more complex basilar dendritic branching. While there was no difference in the extent of branching (**Figure 4L**), FLX neurons had a greater number of 2^nd^ order segments (*p*=.027; **Figure 4M**) and a corresponding increase in nodes on 1^st^ order branches (*p*=.0005; **Figure 4N**). This finding is replicated by an increase in intersections as observed by Sholl analysis extending up to 40 μm away from the cell body (*p*<.05; **Figure S5D**). Other features including dendritic length and dendritic field area were indistinguishable between the two groups at both P10 and P79 (**Table S2**).

In summary, we observed morphological changes to layer V projection neurons in areas related to social behavior deficits. Maternal FLX resulted in less elaborate M1 neurons during development, which persisted into adulthood. The mPFC neurons were more elaborate during development, yet by adulthood this was diminished, presumably due to enhanced dendritic pruning. Together, our morphological data indicate maternal FLX affects dendritic spine formation and arborization in brain regions implicated in social behavior.

### Extended maternal FLX induces repetitive, restricted patterns of behavior

Similar to our analysis of social behaviors, we assessed a range of rodent tasks relevant to repetitive and restricted patterns of behavior to fully characterize the influence of FLX on this symptom domain. In humans, these symptoms can manifest as stereotyped or repetitive motor movements, use of objects, or speech; insistence on sameness, inflexible adherence to routines or patterns; or highly restricted interests. This domain also includes hyper-or hypo-reactivity to sensory input (American Psychiatric Association, 2013). In our mice, we used the marble burying task to examine compulsive digging, spontaneous alternation T-maze to test inflexible adherence to behavior patterns, and Von Frey filaments to gauge reactivity to tactile input. Output from statistical tests for this section is fully reported in **Table 2**.

Mice compulsively dig in bedding, and this behavior is perturbed in models of OCD and ASD (Angoa-Pérez, Kane, Briggs, Francescutti, & Kuhn, 2013), therefore we examined digging in our mice using buried marbles as a proxy. In the *Celf6*-Extended cohort, *Celf6* genotype alone decreased compulsive digging (*p*=.001). In addition, FLX treatment reduced digging in *Celf6*^+/+^ mice (drug × genotype interaction, *p*=.032; *Celf6*^+/+^ mice only, *p*=.0002; **Figure 5A**). However this effect on WT mice did not replicate in the C57-Extended cohort. In the Long Prenatal and Short Prenatal cohorts also, no difference in number of buried marbles was observed (*p*>.05; **Figure 5B-D**). These data suggest that postnatal, but not prenatal, FLX may influence compulsive digging.

**Figure 5.**
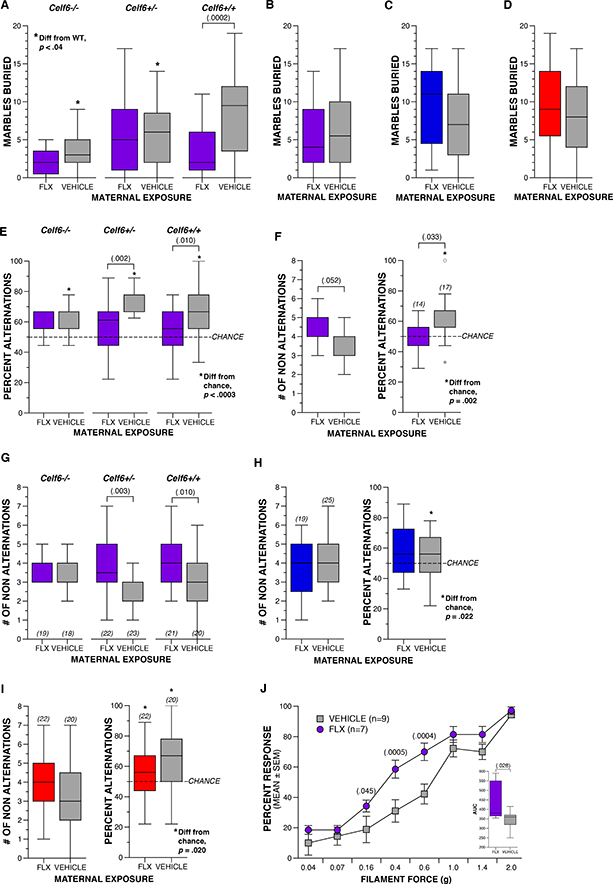
Extended maternal FLX induces repetitive, restricted patterns of behavior. (A) Boxplot of number of marbles buried by *Celf6*-Extended FLX and VEH *Celf6* mutant and WT littermates during adulthood (Genotype × Drug interaction, *p*=.032). (B-D) Boxplot of number of marbles buried by C57-Extended, Long Prenatal, and Short Prenatal FLX and VEH C57BL/6J mice. (E,G) Boxplot of number of non-alternation trials and percent alternation trials in the spontaneous alternation T-maze for Celf6-Extended FLX and VEH *Celf6* mutant and wild type littermates (Drug, *p*=.0001). (F,H,I) Boxplot of number of non-alternation trials and percent alternation trials for C57-Extended, Long Prenatal and Short Prenatal C57BL/6J FLX and VEH mice. (J) Percent of trials during which a response was elicited by Von Frey filament presentation for C57-Extended FLX and VEH C57BL/6J mice (data are means ± SEM; Filament × Drug, *p*=.005). Inset boxplot represents total area under the curve for all filaments per drug group.

In spontaneous alternation on the T-maze, we see differences in the effect of FLX depending on whether exposure was prenatal only or extended. Extended FLX induced perseverative behavior in this task, regardless of genetic background, replicating a percent of alternations better than chance in the VEH mice from Celf6-Extended and C57-Extended cohorts (*p*<.0004 and *p*=.022, respectively; **Figure 5E,F**), but not FLX treated mice. This is also reflected in an increased number of non-alternations in FLX mice (*Celf6*-Extended main effect of drug, *p*=.0001; *Celf6*^+/−^, *p*=.003; *Celf6*^+/+^, *p*=.010; **Figure 5G**, and a trend in the C57-Extended cohort *p*=.052; **Figure 5F**). In contrast, Long and Short Prenatal exposure to FLX did not result in increased non alternation trials (*p*>.05; **Figure 5H,I**). While all exposures induce social behavior deficits, these results suggest that extended exposure is required to induce repetitive or restricted patterns of behavior.

### Maternal FLX results in tactile hypersensitivity

Because we observed abnormalities in marble burying and T-maze performance only in the Extended exposure cohorts, we further examined FLX influence in this cohort on the sensory reactivity aspect of the restricted and repetitive behavior symptom domain. Previously, tactile processing defects were observed in the *Mecp2* and *Gabrb3* models of ASD (Orefice et al., 2016). Therefore we tested tactile sensitivity using the Von Frey filaments in C57-Extended mice and observed hypersensitivity to tactile stimulation: FLX resulted in an increased percent of trials with a response to stimulation compared to VEH mice (*p*=.005; **Figure 5J**) for filaments providing 0.16–0.6g of force (*p*<.046). The area under the curve (AUC) was also greater for FLX compared to VEH mice (*p*=.028), indicating a greater overall response to stimulation across filaments. This tactile hypersensitivity is independent of general activity levels, altered emotionality (anxiogenic behavior), or sensorimotor abilities, as we did not observe differences between Extended FLX and VEH exposure in a 1-hr locomotor activity task (distance traveled, center zone time and entries) or a battery of sensorimotor tasks assessing balance, strength and coordination (data not shown).

### Extended FLX exposure alters stimulus-induced and resting state hemodynamics in mesoscale imaging

OIS imaging, similar to functional MRI, can map brain function and functional connectivity, with use of reflected light rather than magnetic resonance to assess hemodynamics. We tested if an OIS study, leveraging the increased experimental control and genetic homogeneity of a rodent model, can be used to define a mesoscale phenotype mediated by transient SSRI exposure. We adapted a previous pipeline (Bauer et al., 2014) developed for stroke to assess the subtler effects expected for ASD models. To examine both sensory capabilities and baseline brain function we characterized brain activity using both stimulus-evoked response and resting-state functional connectivity (**Figure 6A,B**). Because our first cohort displayed heterogeneity of variance, we imaged a second independent cohort, to assess reliability and robustness of this approach. The second cohort exhibited homogeneity of variance and therefore improved data quality. Analyses of each independent cohort (**Figure S6**) are included below in the description of results.

**Figure 6.**
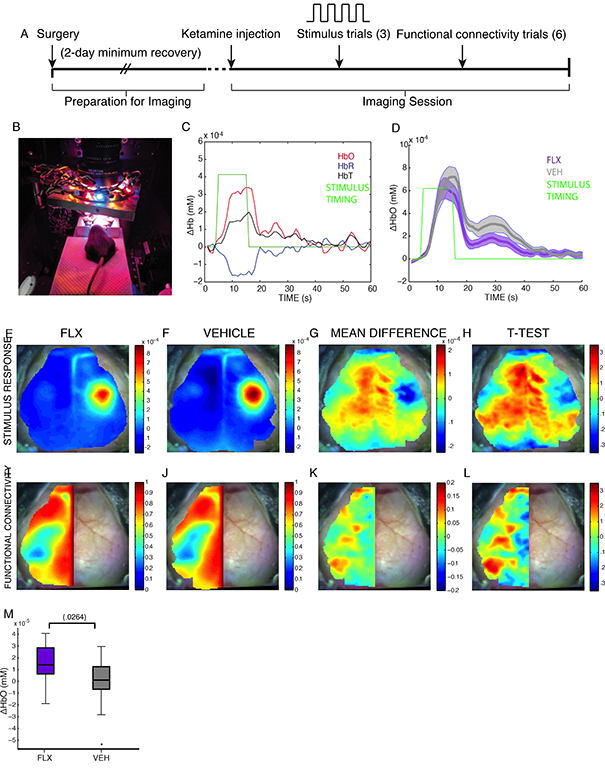
Evoked response to forepaw stimulation and resting-state functional connectivity in FLX mice display temporal and regional HbO response and connectivity differences. (A) Representation of OIS and evoked response imaging workflow. (B) OIS imaging system with mouse: cranial window lies directly below camera and LED ring center, stimulus clips attached to left forepaw under anesthesia. (C) Representative trace of stimulus paradigm. Left forepaw stimulus ( green) was applied at t=5s for a 10-s duration during 5 1-min blocks, averaged to produce traces of HbO change from baseline (red), HbR change (blue), and HbT change (HbO+HbR, black). (D) HbO change during 1-min stimulus block in FLX (purple, n=13) and VEH (gray, n=17) mice (data are means ± SEM). (E-F) HbO activation map at end of stimulus (mean of t=14-16s), averaged between stimulus blocks in each mouse (means ± SEM) between (E) FLX mice and (F) VEH. (G) Mean difference map of FLX-VEH HbO at end of stimulus ( mean of t=14-16s). (H) *t*-statistic map of FLX-VEH HbO at end of stimulus (mean of t=14-16s; uncorrected). (I-J) Contralateral homotopic connectivity maps (mean Pearson correlation coefficient) for (I) FLX and (J) VEH mice. (K) Mean difference map of functional connectivity in FLX-VEH groups. (L) t-statistic map of functional connectivity in FLX-VEH groups (uncorrected). (M) Box plot of global ΔHbO at end of stimulus (mean of t=14-16s).

For the evoked response analysis, left forepaw stimulation was applied to C57-Extended FLX and VEH mice, and changes in oxyhemoglobin (HbO) concentration was observed in the forepaw region of the somatosensory cortex in a block design (**Figure 6C**). HbO response to left forepaw stimulation displayed a faster return to baseline typified by decreased HbO levels in the FLX group during the 45 sec following the end of stimulus (**Figure 6D-H**). This faster recovery to baseline HbO levels replicated in both cohorts of mice (**Figure S6A,D,G**). The AUC of the HbO time trace did not differ significantly between drug groups, although sex x drug interactions were approaching significance (*p*=.090). Pairwise comparison also showed a FLX treatment difference in HbO AUC among males (*p*=.036). The average change from baseline HbO levels at peak stimulation (14-16s) throughout the brain also was higher (*p*=.026) in FLX mice as compared to VEH (**Figure 6M; Video S1,S2**). These data indicate a difference in stimulus-evoked response in the cortex of FLX mice, namely in the FLX group’s faster return to baseline and increased global HbO levels at peak stimulation. This global HbO increase and the lack of a significant difference in amplitude at peak stimulation indicates a more diffuse evoked response in FLX mice.

We next examined FLX mice for alterations in functional connectivity. In the absence of a task or applied stimulus, correlation between individual cortical regions is used to operationally define resting-state functional connectivity independent of structural connectivity (Biswal, Zerrin Yetkin, Haughton, & Hyde, 1995). Specifically, functional connectivity between corresponding regions across hemispheres is high because of the regions’ similar functions, as exemplified by the correlation between left and right motor areas. This homotopic contralateral functional connectivity (HCFC) can be assessed to identify disruption in disease models (Bauer et al., 2014). Comparison of HCFC between FLX and VEH groups (**Figure 6I,J**) identified alterations in the visual cortex (lateral V2; **Figure 6K-L**): FLX mice showed a trend toward increased correlation in the region (*p*=.0330, uncorrected). The HCFC increase in the visual area identified in Cohort 1 replicated in Cohort 2 (**Figure S6M-O**).

These stimulus and functional connectivity data show a difference in evoked response to left forepaw stimulus across time, and identify a region of interest in the visual cortex where increases in HCFC are seen at rest. With these trends replicated between two cohorts, these changes suggest that perinatal FLX exposures also resulted in long term alterations in functional connectivity as assessed by the hemodynamic response in the C57-Extended FLX mice.

## Discussions

Human epidemiological studies suggest SSRI use during pregnancy may increase ASD risk in offspring, however it is unclear from these studies whether ASD susceptibility in offspring is due to maternal psychiatric symptoms or if drug exposure poses an additional risk (Mezzacappa et al., 2017). Here we demonstrate social communication and interaction deficits and repetitive patterns of behavior in offspring of dams exposed to the SSRI FLX during pre-and postnatal development, with corresponding dendritic morphology changes in pertinent brain regions. We further show altered stimulus-evoked cortical response and region-specific decreases in functional connectivity in FLX brains. These findings indicate drug exposure alone is sufficient to induce long-term behavioral, cellular, and hemodynamic-response disruptions in offspring.

Our characterization of ASD-related behaviors included multiple assays in an effort to examine the various social disruptions and repetitive/restricted behaviors observed in autistic individuals. In our mice, maternal FLX induced early social communication deficits, abnormal sociability, and altered social hierarchy behaviors, but did not influence frequency of behaviors observed when an unexposed partner could initiate interactions. We also report increased perseveration and tactile hypersensitivity in FLX pups. Our results suggest FLX exposure spanning perinatal mouse development has the potential to persistently impact social and repetitive behavior. Overall, our findings also show the severity of FLX influence on ASD-related behaviors may depend on developmental period of exposure. Early pregnancy alone was the least vulnerable period of exposure, as we observed increased submissive behaviors in adulthood in mice exposed E0-E16, but control-like early communicative, social approach, and exploration behaviors. FLX exposure through all of gestation influenced early communicative behaviors, sociability, and social hierarchy behaviors and extending exposure through lactation induced perseverative behavior and tactile hypersensitivity. Thus early pregnancy may be the least vulnerable developmental period in the rodent for SSRI exposure, and increasing durations of exposure may increase the extent of behavioral disruptions.

Interestingly, social hierarchy behaviors and repetitive patterns of behavior during adulthood are differentially impacted by prenatal-only and postnatally-extended exposure. A link between low 5HT levels in the mature brain and dominance has been demonstrated in both human and animal research (Kaplan et al., 1994; Uchida et al., 2005). Tryptophan depletion was shown in an adult autistic patient to exacerbate symptoms including perseveration (McDougle et al., 1993), and adult mice fed a tryptophan-depleted diet exhibited increased dominance in the tube test (Uchida et al., 2005). The distinct phenotypes of prenatal-only versus postnatally-extended FLX pups may be mediated by differences in 5HT system development that occurs at these different periods. While 5HT axons reach their targets by birth, terminal field development occurs postnatally (Maddaloni et al., 2017). Excess 5HT during embryonic development acts to down-regulate 5HT innervation through a negative feedback mechanism (Whitaker-Azmitia, 2005) and reduced 5HT terminal processes has also been reported in rodents following postnatal SSRI treatment (Maciag et al., 2006). This suggests prenatal FLX exposure likely influences axonal innervation by 5HT neurons of the raphe, but continued postnatal exposure may have further reduced 5HT terminal fields, possibly meditating the increased dominance and perseverative behavior patterns observed in the Extended FLX cohort.

There is an established body of work in the rodent literature showing clear links between maternal SSRI exposure during pregnancy and a paradoxical increase in depressive-and anxiety-like behaviors in the mature offspring (Avitsur et al., 2016; Boulle et al., 2016; Gobinath, Workman, Chow, Lieblich, & Galea, 2016; Lisboa, Oliveira, Costa, Venâncio, & Moreira, 2007; Noorlander et al., 2008; Olivier et al., 2011; Salari, Fatehi-Gharehlar, Motayagheni, & Homberg, 2016). Our full characterization of ASD-related behavioral consequences across multiple durations of FLX exposure here adds to the limited studies of dam SSRI exposure that have recently begun to focus on these types of behaviors in offspring. Maternal FLX exposure limited to postnatal-only ages (P3-21) was recently shown to not effect social approach behaviors (Nakai et al., 2017) in mice. Together with our findings, this suggests *in utero* exposure is required to influence sociability in adult offspring. Complementing our findings on distinct effects of maternal FLX on dominance, recent work showed prenatal maternal FLX treatment decreased aggressive behaviors, while treatment extending postnatally increased aggressive behaviors in adult C57BL/6 male offspring (Kiryanova, Meunier, Vecchiarelli, Hill, & Dyck, 2016). However, another report showed increased aggression in male offspring of ICR dams exposed to only prenatal FLX (Svirsky, Levy, & Avitsur, 2016). The discrepancies in aggression findings between these two studies may reflect strain × drug interactions. Altogether, rodent research shows clear long-term consequences to SSRI exposure through the dam.

We hypothesized a potentiation of deficits in *Celf6* mutants with FLX exposure due to the effect each manipulation has on the 5HT system. Instead both *Celf6* mutation and FLX exposure independently reduced pup USVs, induced perseveration in the T-maze and reduced digging in the marble burying assay. These complementary behavioral disruptions suggest Celf6 loss and FLX exposure act in a similar manner on the circuits underlying these behaviors, possibly through similar influences on the 5HT system. Interactions between maternal FLX and the 15q11-13 duplication model (15q-dup), which also show reduced brain 5HT levels (Tamada et al., 2010), potentiated deficits in the 15q-dup mice: specifically, hypoactivity and anxiogenic behaviors (Nakai et al., 2017). Maternal FLX actually improved 15q-dup induced sociability, which was linked to restoration of extracellular 5HT levels. We observed a somewhat protective effect of *Celf6* loss in females to FLX-induced sociability deficits. Taken together, these results suggest that in a normal 5HT system, SSRI-induced increase in 5HT activity during development is detrimental to behavioral functioning. However, when the 5HT levels are genetically lowered, SSRI exposure can help restore these levels and prevent the adverse effects on social behavior circuitry. This hypothesis requires further explicit investigation.

Finally, we demonstrate the novel application of OIS to assess stimulus-evoked and functional connectivity in a model of a single ASD risk factor. Human hemodynamic studies in ASD are challenged by the substantial heterogeneity in likely causal mechanisms, making it difficult to determine if there is a strong imaging correlate of ASD. Developing OIS in rodents allows carefully controlled studies of individual risk factors in animals, providing well-powered assessments of the resulting perturbations in mesoscale responses and functional connectivity. In FLX mice, we observed altered cortical hemodynamic response to stimulus. The increased global HbO in FLX mice at stimulation end suggests the evoked HbO response may be more diffuse in FLX mice. While the half-max measure of response is generally localized in somatosensory cortex and does not show a significant difference in peak amplitude, the HbO change from baseline outside this primary region appeared to be stronger in FLX mice. This likely explains the increase in global HbO despite equivalent amplitude to the VEH mice, and indicates a more diffuse hemodynamic response in the FLX mice. The faster return to baseline following stimulation and the trending AUC differences also may reflect the same differences driving the FLX group’s altered tactile sensitivity in the Von Frey assessment. Discrepancies between recovery of excitation versus inhibition have been previously suggested as an explanation for post-adaptation facilitation in somatosensory cortex (Cohen-Kashi Malina, Jubran, Katz, & Lampl, 2013), and the differing shape of the response curve in FLX mice might reflect a similar mechanism. Analysis of the excitatory-inhibitory balance in the cortical response to stimulus in the future may further elucidate the mechanism driving the differences in evoked response across time. While the trend toward increased functional connectivity in the lateral V2 region is not statistically significant, it does suggest that increased power could produce significant findings in functional connectivity in FLX mice and perhaps other models of ASD as well. Application of this imaging pipeline across other rodent models of genetic and environmental risk factors for ASD will allow us to determine if these alterations are a shared consequence of multiple risk factors, and are thus more likely in the causal chain for the behavioral abnormalities in ASD. The same approach may be amenable to testing therapeutic approaches for normalizing this response. Fortunately, OIS in rodents is advancing to allow mesoscale calcium imaging (Wright et al., In Press), which could be used to disambiguate these two possibilities across future studies comparing multiple risk factors.

In spite of a potential for increased risk from FLX exposure, untreated or undertreated depression and anxiety in pregnancy are themselves strongly associated with adverse outcomes (Grote et al., 2010). Thus, while these findings are a contribution to our understanding of ASD pathogenesis, risk and mechanism, as well as our understanding of the developmental role of 5HT, more basic and pre-clinical studies are needed to further characterize the complex effects of SSRI exposure during development.

## METHODS

### Animals

All protocols involving animals were approved by the Institutional Animal Care and Use Committee of Washington University in St. Louis. Mice were house in translucent plastic cages measuring 28.5 cm × 17.5 cm × 12 cm with corncob bedding and standard lab diet and water freely available. The colony room lighting was a 12:12 hr light/dark cycle; room temperature (~20-22°C) and relative humidity (50%) were controlled automatically. All mice used in this study were maintained and bred in the vivarium at Washington University in St. Louis, and were all group-housed. The C57BL/6J wildtype inbred strain and the *Celf6* mutant line were used in this study. Four separate cohorts of mice were used based on maternal drug exposure duration and mouse line: Celf6-Extended, C57-Extended, Long Prenatal and Short Prenatal (**Table S1**) *Celf6* mutant mice were generated on the C57BL/6 background by deletion of exon 4 of the *Celf6* gene as previously described (Dougherty et al., 2013). For the *Celf6*-Extended cohort, heterozygous breedings pairs were used to generate *Celf6*^+/+^, *Celf6*^+/−^, and *Celf6*^−/−^ littermates (**Table S1**). Offspring were genotyped using standard reagents and primers for amplification of the region spanning exons 3 and 4: forward, ATCGTCCGATCCAAGTGAAGC and reverse, CTCCTCGATATGGCCGAAGG. C57BL/6J breeding pairs were used to generate the C57-Extended, Long Prenatal, and Short Prenatal cohorts (**Table S1**). The C57-Extended cohort served to replicate and extend the findings from the *Celf6*-Extended cohort. All mice were examined for USV production, developmental milestones and reflexes, and a subset was used for further behavioral assessment and imaging.

### Maternal SSRI Exposure

In most countries, fluoxetine (Prozac, FLX) was the first SSRI to become available for clinical use (Hiemke & Hartter, 2000). Therefore, FLX is likely to be the most-represented antidepressant in the epidemiology studies of SSRI use during pregnancy and ASD risk. To mimic the 5HT system in human mothers already taking an antidepressant prior to pregnancy, dams were exposed to FLX sat least one week prior to mating. FLX crosses the placental barrier at a rate in mice comparable to that in humans (Noorlander et al., 2008). To avoid inducing unwanted maternal stress that can occur with daily injections, which has been shown to have adverse effects on the developing brain (Matrisciano et al., 2013), FLX was administered orally through drinking water sweetened with 1% saccharin to mask unpleasant drug taste. Control dams received 1% saccharin-only water (VEH). Fluoxetine capsules (20 mg each; Camber Pharmaceuticals, Inc, Piscataway, NJ) were dissolved into water containing 1% saccharin sodium hydrate (Sigma-Aldrich). The FLX dose used in this study was equivalent to the maximum recommended human dose (MRHD) of 80 mg per day on a mg/m^2^ basis (Marken & Munro, 2000). The dose calculations are based on equivalent surface area dosage conversion factors (Freireich, Gehan, Rall, Schmidt, & Skipper, 1966) and approximate drinking water consumed daily (Bachmanov, Reed, Beauchamp, & Tordoff, 2002). Average drug water intake per day was recorded throughout the study to monitor drug exposure levels. The FLX water was prepared so that each mouse would consume 48 mg/day (16 mg/kg/day based on a 30 g mouse) or 6.5 mL/day of 0.074 mg/mL FLX in 1% saccharin water. Females of the same drug group were co-housed to reduce stress induced by isolation housing, and placed into a the cage of a single-housed male for breeding. Upon detection of a vaginal plug following breeding, the females were removed from the male to isolate maternal drug exposure effects and avoid paternal drug exposure. Three drug exposure durations were used. Extended exposure continued until P14, the age just before pups begin to consume food and water, to avoid direct drug exposure in the pups. Long Prenatal exposure lasted until birth of the pups, and Short Prenatal exposure was stopped at E16 (**Figure 1A**).

### Behavioral Tasks

Multiple behavioral assays across the same domain were employed to adequately determine presence of behavioral disruptions. Experimenters were blinded to experimental group designations during behavioral testing. Order of and age at testing were chosen to minimize effects of stress and previous testing. Developmental reflexes and milestones assessment of the *Celf6*-Extended, Long Prenatal, and Short Prenatal cohorts occurred on P5 – P14. Adult behavioral testing for all cohorts began at P60. Adult behavioral testing of the *Celf6*-Extended cohort included a battery of sensorimotor measures, followed by the social approach test, marble burying, spontaneous alternation T-maze, and the 1-hr locomotor activity task. Mice in the C57-Extended cohort were assessed in the juvenile play task P22 – P30, and adult behavioral testing included marble burying, spontaneous alternation T-maze, the tube test of social dominance, and the Von Frey assessment of tactile sensitivity. Both the Long Prenatal and Short Prenatal cohorts were tested as adults in the social approach test, followed by spontaneous alternation T-maze, marble burying, and the tube test of social dominance.

#### Maternal isolation-induced USVrecording

Ultrasonic vocalizations (USVs) are considered a strongly conserved affective and communicative display that elicits maternal search and retrieval responses, nursing, and caretaking, and is used in the rodent literature to model early communicative deficits (Haack, Markl, & Ehret, 1983). Playback experiments demonstrated lactating dams respond rapidly with searching behavior to pup isolation calls. In addition, these dam behaviors are dependent on acoustic call features, such as duration and frequency, suggesting these features have communicative value (Wöhr et al., 2008). USV production due to maternal isolation in the C57BL/6J mouse pup normally peaks just after P7, disappearing completely by P14 (Rieger & Dougherty, 2016).

For this study, USV recording occurred on P5, 7, and 9. Dams were removed from the home cage and placed into a clean standard mouse cage for the duration of testing. Pups in the home cage were placed into a warming box (Harvard Apparatus) for at least 10 minutes prior to the start of testing to control temperature. Skin surface temperature was recorded immediately prior to placement in the USV recording chamber via a noncontact HDE Infrared Thermometer during ensure consistent temperatures as lower body temperature of the pup is known to increase USV production (I. Branchi, Santucci, Vitale, & Alleva, 1998). Differences in temperature between FLX and VEH pups were not detected, indicating the differences in USV production were not secondary to thermoregulation differences. For recording, pups were individually removed from the home cage and placed into an empty standard mouse cage (28.5 cm × 17.5 cm × 12 cm) inside a sound-attenuating chamber (Med Associates, Saint Albans, VT). USVs were obtained using an Avisoft UltraSoundGate CM16 microphone, Avisoft UltraSoundGate 416H amplifier, and Avisoft Recorder software (gain=2 dB, 16 bits, sampling rate=250 kHz). Pups were recorded for 3 min, after which they were weighed and returned to their home cages inside the warming box. Tissue from a toe was also collected at this time on P5 for genotyping. Frequency sonograms were prepared from recordings in MATLAB (frequency range=25 kHz to 120 kHz, FFT size=512, overla*p*=50%, time resolution=1.024 ms, frequency resolution=488.2 Hz), and individual syllables and other spectral features were identified and counted from the sonograms according to a previously published method (Maloney et al., In Press).

#### Developmental reflexes and milestones assessment

Mice were evaluated at several time points for achievement of physical and behavioral milestones of development. A visual check for the presence of detached pinnae was done at P5, and eye opening at P14. Weight was measured at P5, 7, 9 and 14, concurrent with USV recordings and righting reflex testing. To assess surface righting reflex at P14, each mouse was placed in a 50 ml conical containing a lid with a hole. When the belly of the mouse was facing down, the conical was quickly turned 180 degrees in a smooth motion placing the mouse on its back. The time for the mouse to right itself with all 4 paws underneath its belly was recorded up to 60 sec. Each mouse received 3 trials, which were averaged for analysis.

#### Juvenile play social interaction

Full-contact social behaviors were assessed through juvenile play interactions using a procedure adapted from previously published methods (Peñagarikano et al., 2011). Mice were tested between P22 – P30, and were paired with an age-and sex-matched C57BL/6J stimulus mouse derived from standard mouse breeding. All mice were weighed prior to testing. The procedure consisted of 3 consecutive 10 min trials. During trial 1, the stimulus mouse was habituated to the testing chamber. For trial 2, the test animal was habituated to the chamber while the stimulus mouse was placed in a holding chamber lined with clean corn cob bedding. For the third trial, the stimulus mouse was placed back into the testing chamber with the test mouse and their interactions were recorded for 10 minutes. The testing chamber was cleaned with 70% ethanol between test animals and the corn cob bedding was replaced. The test apparatus was a transparent enclosure (25 × 15 × 12 cm) containing a layer of clean corn cob bedding on the floor and surrounded by a clear acrylic enclosure measuring 28 × 17.5 × 37.5 cm. A 4 cm diameter hole on the top of the enclosure allowed for placement of a digital video camera (Sony HDR-C×560V High Definition HandyCam camcorder) to record scenes inside the apparatus. The apparatus was housed inside a custom built sound-attenuating chamber (70.5 × 50.5 × 60 cm), which was equipped with two LED infrared lights (Crazy Cart 48-LED CCTV infrared Night Vision Illuminator) to allow for capture of social behaviors in darkness.

Video files in MPG format were acquired in 360 × 240 or 544 × 362 pixel resolution with a frame rate of 25 or 30 frames per second. Videos were minimally post-processed to key only grayscale images, remove associated audio track, and convert to AVI containers before tracking. Simultaneous supervised tracking of both the stimulus and experimental animals was performed in MiceProfiler (de Chaumont et al., 2012) on the Icy platform, with scale value of 0.35 and pixel intensity threshold used to identify mice optimized for each video as necessary to ensure most accurate tracking. Manual corrections of tracking was performed as necessary through the course of each video. Two videos were excluded due to unexpected differences in zoom and resolution and eleven other videos were excluded because one mouse left the field of view for a portion of the ten minute testing time.

Tracked videos were then processed using a custom pipeline in MATLAB as follows. MiceProfiler data points for each frame and <x,y> positions of head, center of mass (“body”), and tail were parsed from the XML tracking data, pixel coordinates were converted to centimeters using the real world size of the testing apparatus, and frame number converted to time in seconds using the frame rate. Occasionally isolated frames contained missing data points occur where MiceProfiler does not record a value, and these were recorded as NaN (not-a-number) in MATLAB. Because of these occasional missing values, and jitter which occurs during tracking, data were smoothed using a 11-point moving average smooth. This smooth when rewatched within MiceProfiler helped to ensure more accurate tracking. After smoothing, positional values for head, body, and tail were used to estimate two-dimensional kinematics, using the first difference approximation for derivatives: velocity, acceleration, and jerk. Vectors defined by the head and tail positions were used to determine relative orientation of the two mice in the field of view, and final processed data contained the following variables by frame: distance traveled, length of body axis (head-to-tail) and direction (radians) with respect to the field of view (coordinate system <0,0> in lower left), the direction (radians) and magnitude of each 2D component of motion (velocity, acceleration, jerk) for each animal, and inter-animal parameters (angle between both animals and between their velocity vectors, all pairwise distances in cm between head, body, tail), from which total distance traveled and average speed (cm/s) were determined. Thresholds of 3.502 cm for head-to-head distance and 3.125 or 3.145 cm head-to-tail distance were used to define head sniffing and anogenital sniffing behaviors, respectively. These thresholds were determined through examination of the histogram of all head-to-head and head-to-tail distances across all videos and verified by manual inspection of video after applying threshold. After thresholding, bouts of behavior were scored as frames with distances below threshold, and bouts separated by 35 frames or less (<= 0.10 seconds or <= 0.17 seconds) were merged. From these, fraction of total frames for each behavior, as well as number and average duration of bouts of behavior were determined. Measures of overall activity per mouse, such as distance traveled and average speed, were also extracted.

#### Social approach

The social approach task was used to quantify sociability and preference for social novelty, and as previously described (Moy et al., 2004). Sociability was defined here as a tendency to pursue social contact. Preference for social novelty was defined as pursuing social contact with a novel conspecific as compared to a conspecific from a previous interaction. The social approach testing apparatus was a rectangular clear acrylic box divided into three separate chambers each measuring 19.5 cm × 39 cm × 22 cm including clear acrylic dividing walls with rectangular openings measuring 5 cm × 8 cm to allow for movement between chambers, which could be shut off by sliding down clear acrylic doors. This clear acrylic apparatus was housed inside a custom built sound-attenuating chamber (70.5 × 50.5 × 60 cm), lit with LED Flex Ribbon Lights (Commercial Electric, Home Depot, Atlanta, GA) with to provide approximately 20 lux illumination in the chamber. A small stainless steel conspecific cage (Galaxy Pencil/Utility Cup, Spectrum Diversified Designs, Inc, Streetsboro, OH), measuring 10 cm in height and 10 cm in diameter at its base, was placed in each outer chamber, and had vertical bars that allowed minimal contact while preventing fighting. A CCTV camera (SuperCircuits, Austin, TX) connected to a PC computer running the software program ANY-maze (Stoelting Co., Wood Dale, IL) tracked the movement of the mouse within the apparatus and time spent in each investigation zone surrounding the conspecific cages. The investigation zones encompassed an area of 2 cm around the conspecific cages. Only the head was tracked in the investigation zone to quantify intention to investigate the conspecific. Total distance traveled was also ascertained as an index of general activity levels. The entire apparatus was cleaned between animals with a chlorohexidine diacetate solution (Nolvasan, Zoetis, Parsippany-Troy Hills, NJ). The conspecific cages were cleaned with 70% ethanol solution between each mouse.

The social approach task consisted of four, consecutive 10 min trials. For the first trial, the mouse was placed in the middle chamber with the doors to the outer chambers shut and allowed 10 min to habituate to the apparatus. During the second trial (habituation trial), the mouse was allowed to freely investigate and habituate to all three chambers for 10 min. Performance of the mouse during the third trial (sociability trial) allowed for the evaluation of sociability to an unfamiliar, sex-matched conspecific (C57BL/6J) placed in one conspecific cage versus an empty conspecific cage. Again, the mouse was allowed to move freely within the apparatus for 10 min. During the fourth trial (preference for social novelty trial), the now familiar conspecific remained in the apparatus, and a new, unfamiliar sex-matched conspecific (C57BL/6J) was placed in the other conspecific cage. The mouse was allowed to move freely within the apparatus for 10 min, and the mouse’s preference for social novelty was quantified. Placement of conspecifics was counterbalanced.

#### Tube test of social dominance

Under laboratory conditions, mice begin to develop social hierarchy behaviors at 6 weeks of age, which result in dominance ranks within their social groups (Hayashi, 1993). The tube test of social dominance allows for examination of social dominance rank between two pairs of mice after 8 weeks of age, and was adapted from previously described methods (Wang et al., 2011). The apparatus consisted of a clear acrylic tube measuring 3.6 cm in diameter and 30 cm in length. This task spanned five consecutive days. On days 1 and 2, each mouse was exposed to the test apparatus to habituate the animals to the testing tube and to walking through the testing tube to the other side. This was conducted from each side of the tube. On days 3 – 5, dominance bouts were conducted with sex-matched pairs of FLX and VEH mice, avoiding cage mate pairings. A new pair was used for each bout such that each mouse was paired with three distinct partners, and side of entry was alternated. For each bout, a small acrylic divider was placed in the center of the tube, prohibiting the animals from crossing the center, and each mice was allowed to enter the tube from one end. Once the animals met in the center, the divider was lifted and the bout lasted 2 minutes or until one animal was backed out of the tube by the other (all four paws exiting the tube). The animal remaining in the tube was the winner of the bout (dominant) and the animal that was backed out was the loser of the bout (subordinate). The bouts were recorded with a USB camera connected to a PC laptop (Lenovo) and subsequently scored by an observer. The percent of bouts won was calculated for each mouse, and compared between groups. The acrylic tube was cleaned with a chlorohexidine diacetate solution (Nolvasan, Zoetis, Parsippany-Troy Hills, NJ) between each bout. On each day, male bouts were conducted first, followed by female bouts.

#### Marble burying task

Marble burying behavior in mice serves as a proxy for repetitive and perseverative digging behavior (Angoa-Pérez et al., 2013), and our procedure was adapted from these previously described methods. The apparatus was a transparent enclosure (47.6 × 25.4 × 20.6 cm) housed within a sound-attenuating chamber (70.5 × 50.5 × 60 cm), lit with LED Flex Ribbon Lights (Commercial Electric, Home Depot, Atlanta, GA) to provide approximately 20 lux illumination. Each enclosure was filled with 3 cm of clean, autoclaved corncob bedding. Using a template, 20 clear marbles were placed in 5 rows of 4. For testing, the mouse was placed in the center of the enclosure, and allowed to freely explore for 30 minutes. The animal was then removed and two independent observers scored buried marbles. A marble was considered buried when at last 2/3 of it was covered by bedding. The average score between the two observers was used for analysis. The correlation between observers’ scores for all marble burying experiments in this study was *r* > .92, *p*=.000. In between animals, new fresh, autoclaved bedding was used and all marbles were cleaned thoroughly with 70% ethanol.

#### Spontaneous alternation T-maze

The spontaneous alternation T-Maze was used to assess perseverative exploratory behavior and was adapted from previously published methods (Peñagarikano et al., 2011). Testing was conducted under dim overhead lighting. The apparatus was made of opaque acrylic and comprises a 20 × 8.7 cm start chamber with two radiating arms, each measuring 25 × 8.7 cm. Removable doors were used to sequester the animal in the start box, or either maze arm. Testing consisted of 10 consecutive trials, each lasted 2 minutes or until the animal made an arm choice. For each the first trial, the animal was placed in the start box with the door closed for two minutes to habituate to the apparatus. The door was then removed and the animal allowed to explore either the right or left arm of the maze. An arm choice was determined when the animal entered the arm with all four paws. Then the door to that arm was closed, and the animal allowed to explore it for 5 seconds. The door was again lifted and the animal was allowed to return to the start box and the door shut. If the animal did not quickly move back to the start area, it was gently guided by placement of a hand or object behind the animal, yet avoiding picking the animal up by the tail and moving back to the start chamber as this can result in a negative association with that arm and impact the spontaneous alternation. After 5 seconds, the start box door was again lifted to start the next trial. If no arm choice was made after 2 minutes, the animal was gently guided back to the start box. After 10 consecutive trials, the animal was returned to its home cage and the apparatus cleaned thoroughly with a chlorohexidine diacetate solution (Nolvasan, Zoetis, Parsippany-Troy Hills, NJ). Each of the two trials was scored as an alternation, a non-alternation or no choice trial. The number of non-alternations and percent of trials alternating were compard between groups.

#### Von Frey assessment of tactile sensitivity

The Von Frey task assessed reflexive, mechanical sensitivity to a punctate stimulus, and was conducted as previously described (Mickle, Shepherd, Loo, & Mohapatra, 2015). The testing apparatus consisted of a metal grid surface elevated 63.5 cm, which allowed access to the plantar surface of the animals’ paws. On top set individual acrylic boxes (10 cm × 10 cm × 10 cm) open on the bottom and opaque on three sides to prevent visual cues between animals. All mice were acclimated to the testing room 30 min prior to habituation and testing. On days 1 and 2, all mice were habituated to the testing apparatus for 1 hour. On day 3, mice were allowed to acclimate to the testing apparatus for 30 minutes prior to start of testing. Eight different Von Frey hair filaments (0.04 – 2 g; Stoelting Co., Wood Dale, IL) were applied to the plantar surface of each animal’s hind paw and withdrawal responses were recorded. Presentations started with the lowest filament strength (0.04 g) and increased to the maximum filament strength (2 g). Each filament was applied to the plantar surface of each hind paw five times, and the number of paw withdrawal responses was recorded as percentage of responses. To evaluate the changes in paw withdrawal responses to the whole range of filaments over the testing duration, the area of the curve (AUC) was calculated for each animal.

#### 1-hr locomotor activity

A 1 hr locomotor activity/exploration test was conducted to assess the general activity, exploratory behavior, and emotionality of the mice adapted from previously published methods (Dougherty et al., 2013). This test also served as a control test to identify any differences in general activity that may interfere with the interpretation of cognitive, social, and/or emotionality tests. The mice were evaluated over a 1 hr period in transparent enclosures (47.6 × 25.4 × 20.6 cm). A digital video camera connected to a PC computer running ANY-maze (Stoelting Co., Wood Dale, IL.) tracked the movement of the animal within a 33 × 11 cm central zone and a bordering 5.5 cm peripheral zone. General activity variables (distance traveled and time at rest) along with measures of emotionality, including ‘time spent’, ‘distance traveled’ and ‘entries made into the central zone’, as well as ‘distance traveled in the peripheral zone’ were analyzed. Each enclosure was cleaned with 70% ethanol solution between each mouse.

#### Sensorimotor battery

Balance, strength, and coordination were evaluated by a battery of sensorimotor measures using previously published procedures (Dougherty et al., 2013).The battery included walking initiation, ledge, platform, pole, and inclined and inverted screen tests. An observer manually recorded time in to hundredths of a second using a stopwatch for each test. Two trials were conducted for each test and the average of the two yielded a single time, which was used in the analyses. To avoid exhaustion effects, the order of the tests during the first set of trials was reversed for the second set of trials. The order of the tests was not counterbalanced between animals so that every animal experienced each test under the same conditions. All tests lasted a maximum of 60 s, except for the pole test, which lasted a maximum of 120 s. The tests are described below.

The walking initiation test assessed the time taken by a mouse to move out of a small area. The mouse was placed on a flat surface inside a square measuring 21 cm × 21 cm, marked on the surface of a supply cart with white tape. The time for the mouse to leave the square was recorded, i.e. all four limbs concurrently outside of the square. Basic balance ability was assessed by the performance on the ledge and platform tests. The ledge test required the mouse to balance on a clear acrylic ledge, measuring 0.50 cm wide and standing 37.5 cm high. Time the mouse remained on the ledge was recorded. During the platform test, the mouse used basic balance ability to remain on a wooden platform measuring 1.0 cm thick and 3.3 cm in diameter and elevated 27 cm above the floor. The time the mouse was able to balance on the platform was recorded. The pole test was used to evaluate fine motor coordination. The mouse was placed head upward on a vertical pole with a finely textured surface and the time taken by the mouse to turn downward 180° and climb to the bottom of the pole was recorded. The 60°, 90°, and inverted screen tests assessed a combination of coordination and strength. The mouse was placed head oriented downward in the middle of a mesh wire grid measuring16 squares per 10 cm, elevated 47 cm and inclined to 60° or 90°. The time required by the mouse to turn upward 180° and climb to the top of the screen was recorded. For the inverted screen test, the mouse was placed head oriented downward in the middle of a mesh wire grid measuring16 squares per 10 cm, elevated 47 cm, and, when it was determined the mouse has a proper grip on the screen, it was inverted to 180°. The time the mouse was able to hold on to the screen without falling off was recorded.

### Golgi-Cox Impregnation and Morphological Analysis

The Golgi-Cox method (Lee & Lee, 2012) was used to quantify dendritic morphology. This technique, while almost 150 years old, remains a standard histological technique for its ability to stain neurons in their entirety, including cell bodies, axons, dendritic processes, and synaptic spines. Importantly, Golgi-Cox stains approximately 1-2% of neurons present, resulting in clearly distinguishable neurons and processes, allowing for stereological tracing of the dendritic arbors of single neurons (Das, Reuhl, & Zhou, 2013).

#### Tissue Preparation

Offspring from FLX or VEH dams were deeply anesthetized using isoflurane and then euthanized via decapitation at either P10, a developmental age at which we observed USV deficits, or P79, an age at which increased social dominance behaviors were observed (**Figure 3A**). Brains were rapidly dissected and stained via Golgi-Cox impregnation using the FD Rapid GolgiStain kit (FD NeuroTechnologies, INC, Ellicott City, MD). All procedures followed the manufacturer’s manual with minor deviations as reported below. Freshly dissected brains were rinsed with Milli-Q water and immersed in impregnation solution (equal parts A solution and B solution) mixed 24 hours previously and stored in the dark. Impregnation solution was replaced with fresh solution after the first 6 hours. Incubation lasted 21 days for P10 brains and 7 days for P79 brains, with continuous agitation. Brains were transferred to Solution C for 72 hours, followed by flash freezing via immersion into-70°C isopentane, then were stored at -80°C until sectioning. Tissue was coronally sectioned at 200 μm using a Leica CM1950 cryostat and immediately mounted on gelatin-coated slides using a small drop of Solution C. Within 72 hours, slide-mounted sections were stained as outlined in the manufacturer’s manual. Slides were rinsed 2 times for 4 min each in Milli-Q water, then placed in 1:1:2 mixture Solution D:Solution E: Milli-Q water for 10 min. Sections were again rinsed 2 times for 4 min each in Milli-Q water, followed by dehydration in 50%, 70% and then 95% ethanol for 4 min each and then 100% ethanol 4 times for 4 min each. Sections were cleared in Citrasolv and coverslipped using Permount and allowed to dry for two weeks in the dark at 4°C before microscopy.

#### Neuron Selection, Morphological Quantification

For each region, age, and drug condition, at least 10 layer V pyramidal neurons were selected for quantitative analysis of dendritic branching and spine densities. The selected neurons had dendritic arbors that were fully impregnated with stain and were not truncated from slicing. The apical dendrite tracings were often truncated, therefore only morphological features of the basilar dendrites were quantified. Each neuron had several spiny basilar dendrites extending from the cell body and a single apical dendrite (**Figure 3C, 4B,J**). The selection area of M1 was based on the location active during USV, as described by Arriaga et al. (Arriaga et al., 2012) (between −0.02 mm and +0.40 mm from bregma; **Figure 3B**). At this coronal level, the anterior commissure bridges the hemispheres at +0.13 mm (Franklin & Paxinos, 2008), therefore neurons were traced that occur in the section where the anterior commissure bridges +/− one section anterior or posterior, where the limbs of the anterior commissure are well-defined but do not bridge the hemispheres. The sampled area of the mPFC (about +1.94 mm from bregma; **Figure 4A**) was motivated by studies of Wang et al. (Wang et al., 2011), showing that the synaptic efficacy of neurons in this area of mPFC mediated social dominance. This sampled region of mPFC is posterior to the olfactory bulb, anterior to the the basal ganglia, and contains the most anterior extent of the corpus callosum (anterior forceps) and the anterior commissure (Franklin & Paxinos, 2008), and corresponded to the anterior cingulate cortex, prelimbic area and infralimbic area (Wang et al., 2011).

All neurons were manually traced using a 60× oil objective on a Nikon E800 Eclipse microscope and Microfire A/R camera by Optronics with a Nikon Optem 1X DC10NN adapter connected to a Dell Precision T3600 computer equipped with Neurolucida software (MBF Bioscience, Williston, VT). Based on our previously published methods, tracing involved drawing the maximum contour of the soma, following all dendrites along their entire length, and marking all visible dendritic spines (Bianchi et al., 2011). Neuronal morphology was quantified according to eight measures adapted from Bianchi et al. (Bianchi et al., 2011) including number of dendritic spines, total dendritic length, dendritic spine density (ratio of spines per 1 μm dendritic length), number of branching segments, number of branching nodes, number of branching segment endings, total dendritic field area (total area encompassed by the branched structure; μm^2^), soma area (μm^2^) and highest branching order. Branch order is defined by level removed from soma, such that the dendritic branches originating from the soma are considered first-order branch segments, and branches originating from first-order are second-order segments, and so on. As most apical dendrites passed out of plane of sectioning, only basilar dendrites were included in quantitative analyses. In addition to these measures a Sholl analysis was performed, which assessed neuronal complexity as it spatially relates to the soma. In short, the Sholl analysis involved placing concentric circles with gradually increasing radii centered at the centroid of the soma. For each neuron, we tracked the number of times dendrites intersect with a circle of a given radius.

All tracings were obtained by two researchers (C.J. and S.A.), who were normed to an experienced rater (A.L.B.) (Bianchi et al., 2011) and checked by the primary investigator (S.E.M.). As assessed by the coefficient of intraclass correlation, interrater reliability was high in all measures of interest (C.J.-S.A.): dendritic spines=.817; dendritic length=.968; branching nodes=.885; branch segment endings=.956; and soma area=.805.

### Optical Instrinsic Signal (OIS) Imaging

#### Animal Preparation

Adult C57Bl/6J mice (12-18 weeks of age) from the C57-Extended cohort were used for imaging and processed in two independent cohorts (cohort 1: FLX n=7, VEH n=8; cohort 2: FLX n=7, VEH, n=9). These mice were a subset of those used for the behavior testing described previously. Mice were fitted with a transparent Plexiglas window to facilitate imaging. Mice were sedated using isoflurane (3% induction, 1% maintenance, 0.5 L/min) with body temperature maintained at 37°C using a heating pad; the head was then shaved and the mouse prepared for surgery in a stereotactic restraint. An incision in the scalp was made along midline to expose the skull, and the skin was reflected to expose an approximately 1 cm^2^ cortical field of view through the skull. A cranial window of Plexiglas was affixed to the dorsal surface of the skull using dental cement (C&B-Metabond, Parkell Inc., Edgewood, NY, USA) allowing chronic imaging.

After a minimum of 48 hours of post-surgery recovery, the mice were anesthetized for imaging by intraperitoneal injection of ketamine/xylosine cocktail (86.9 mg/kg ketamine, 13.4 mg/kg xylazine, dosage 0.005 mL/kg), and maintained at 37°C by heating pad (mTCII, Cell Microcontrols). The mouse’s head and imaging plane were secured using a platform mount which attached to and stabilized the Plexiglas window adhered to the mouse’s skull.

#### Imaging System

Sequential illumination was provided at four wavelengths by a ring of light-emitting diodes (LEDS: 478 nm, 588 nm, 610 nm, and 625 nm; RLS-5B475-S, B5B-4343-TY, B5B435-30S, and OSCR5111A-WY, respectively, Roithner Lasertehnik) approximately 10 cm above the mouse’s head. Diffuse reflected light was captured by a cooled, frame-transfer EMCCD camera (iXon 897, Andor Technologies). Acquisition was synchronized with the LEDs’ sequential illumination and controlled via custom-written software (MATLAB, Mathworks) at a full frame rate of 30 Hz. Field of view was approximately 1 cm^2^ with an anterior-posterior view from olfactory bulb to superior colliculus, consisting of 512 × 512 pixels with pixel size of approximately 80 μm × 80 μm at data analysis. Imaging protocols and processing are modified from those described previously (Bauer et al., 2014).

#### Functional Connectivity

Functional connectivity data under anesthesia and without stimulation (referred to as resting state here) was collected in each mouse for up to 30 minutes, split into 6 consecutive runs of 5 minutes. Visual monitoring of the mouse was performed throughout, and upon the detection of whisking or other movement indicating a return to wakefulness, the imaging session was terminated and the mouse removed from the imaging system to complete its emergence from the anesthetized state.

#### Forepaw Stimulation

Electrical pulses were generated by an isolated pulse stimulator (Modell 2100, A-M Systems, Carlsborg, WA, USA) and administered to the left forepaw by micro vascular clips (Roboz Surgical Instrument Co., Gaithersburg, MD, USA). Stimulation data was collected in 3 consecutive runs of 5 minutes each. The 5-minute run consisted of 5X 1-minute blocks; each with an initial 5s rest period, followed by 10s of electrical stimulation (frequency: 3 Hz, pulse duration: 300 μs, current: 0.75 mA), and 40s of rest. The 5X 1-minute blocks were later block averaged for each stimulus trial.

#### Imaging Data Processing and Analysis

Data were processed using MATLAB and the imaging analysis pipeline previously described in literature (Bauer et al., 2014) with modifications as described. Image light intensity was interpreted using the Modified Beer-Lambert Law, and absorption coefficient data were converted to hemoglobin concentration changes. Each pixel’s time series was downsampled from 30 Hz to 1 Hz, and analysis was performed on pixels corresponding to brain tissue only, as determined by a binary brain mask made using a false white light image produced from the collected data. Image coregistration was performed by affine-transforming images to a common atlas space. The junction between coronal and sagittal sutures directly posterior to the olfactory bulb and lambda were used as landmarks to standardize seed placements and brain regions across mice. A data quality check was performed on each run’s data, at which point runs were discarded which showed significant noise compared to signal, namely when reflected light intensity varied by 1% or greater for any of the wavelengths, movement of the animal was detected by pixel displacement during the run, or stimulus application failed and no change from baseline was registered. The runs which passed this quality-control step were used to create average functional connectivity maps or stimulus time traces for each mouse.

The processed resting-state functional connectivity and evoked-response data produced were used to analyze HCFC and ΔHb across time, respectively. Global signal regression was performed on all data to remove sources of variance, except where indicated otherwise. An averaged map of pixel-by-pixel comparisons of HCFC for each mouse was produced by use of the brain mask and atlas-based midline coordinates. The Paxinos atlas (Franklin & Paxinos, 2008) was used for all HCFC regional comparisons, in which all pixels within the masked atlas region were averaged between mice and within region to produce a single Pearson correlation coefficient per region.

Forepaw stimulation data were analyzed both across time and at stimulus end (mean across t=14-16s). Each stimulus block was normalized to average baseline hemoglobin levels at t=0s to t=5s, the rest period prior to stimulus onset. To calculate change in hemoglobin levels, the time traces of all pixels at half-max at the mean of t=14-16s were averaged and plotted across time. Measurements in ΔHbO_2_ were used to represent change in primary figures (**Figure 6D-M**), although ΔHb_T_ (total hemoglobin change) can also be used to represent activation changes. ΔHbT measures demonstrated a similar time trace when calculated (**Figure S6C,F,I**). Pearson *r* values were Fisher z-transformed prior to all statistical comparisons.

## QUANTIFICATION AND STATISTICAL ANALYSIS

All statistical analyses were performed using the IBM SPSS Statistic software (v.24) except where otherwise stated. Prior to analyses, all data was screened for missing values, fit between distributions and the assumptions of univariate analysis, and homogeneity of variance. Analysis of variance (ANOVA), including repeated measures ANOVA, or mixed models were used to analyze the behavioral data where appropriate, with main factors of sex and drug exposure The Huynh-Feldt adjustment was used to protect against violations of sphericity/compound symmetry assumptions where appropriate. Multiple pairwise comparisons were subjected to Bonferroni correction when appropriate. Chi-square goodness of fit test was used to assess categorical variables. Differences between morphometric features of Golgi-Cox staining were determined with t-tests and appropriate corrections. For OIS analyses, statistical significance was determined by unpaired, two-tailed t-tests assuming unequal variance in MATLAB. Tukey’s HSD or the Games-Howell method were used as post hoc tests. Probability value for all analyses was *p*<.05 except where otherwise stated. Test statistics and other details for each experiment are provided in Tables 1 and 2.

## Acknowledgments

We would like to thank Andreas H. Burkhalter, PhD, Rinaldo D’Souza, PhD, Durga P. Mohapatra, PhD, Karen O’Malley, PhD and Steven Harmon for access to equipment and training, as well as Matthew Reisman for advice and discussion, and Rayden Hollis for equipment construction.

This work was funded by W.M. Keck Fellowship in Molecular Medicine from Washington University in St. Louis to SEM, the McDonnell Center for Systems Neurosciences (JDD, JC), and the NIH (5U01MH109133-02, 5R00NS067239), and the JDD is a NARSAD Independent Investigator. Authors have no conflicts of interest to disclose.

